# DRUMBEAT: Temporally resolved interpretable machine learning model for characterizing state transitions in protein dynamics

**DOI:** 10.1101/2025.08.04.668534

**Authors:** Babgen Manookian, Elizaveta Mukhaleva, Grigoriy Gogoshin, Supriyo Bhattacharya, Nagarajan Vaidehi, Andrei S. Rodin, Sergio Branciamore

## Abstract

Conformational transitions are central to protein function, yet their mechanistic analysis remains challenging due to the multi-dimensionality and timescales underlying the molecular motions. While interpretable network models such as Bayesian networks have advanced the identification of key residue interactions in molecular dynamics (MD) data, they lack temporal resolution and cannot capture the sequence of events during transitions. Here, we introduce **D**ynamically **R**esolved **U**niversal **M**odel for **B**ay**E**si**A**n network **T**racking or DRUMBEAT, a machine learning approach that combines a universal graph topology with sliding-window rescoring to generate interpretable, time-resolved maps of cooperative events in MD trajectories. Applying DRUMBEAT to the benchmark Fip35 WW domain folding trajectories from DE Shaw Research Group, we recover both major folding pathways and critical residues previously highlighted by experiment. Importantly, DRUMBEAT provides new insight in two ways: (1) uncover unknown protein features important for transition, and (2) dissect the order and timing of conformational changes, revealing the precise sequence of residue contact closures during individual folding events. Robustness analysis demonstrates that both the universal graph and time-resolved results are highly consistent across multiple sampling replicates. These findings establish DRUMBEAT as a scalable and interpretable machine learning framework for dissecting the dynamics of protein folding and other conformational transitions, offering a generalizable tool for the mechanistic study of biomolecular dynamics.

## Introduction

Proteins exist as dynamic ensembles, continually interconverting among conformational states on timescales from picoseconds to milliseconds. Transitions between these states underpin fundamental biological processes, including enzyme catalysis, signal transduction, and ligand binding. Kinetic studies have linked catalytic turnover rates to microsecond-scale motions in active-site loops^1,2^, while NMR relaxation and single-molecule FRET have revealed that allosteric regulation often involves transient, low-population states connecting distant functional sites.^3,4^ The presence, lifetime, and ordering of such intermediates determine reaction efficiency and specificity, including drug-binding selectivity, thus making accurate analysis of their emergence and decay essential for mechanistic biochemistry and structure-based design.

Pairwise-correlation and contact-frequency graphs are widely used to visualize communication pathways in molecular dynamics (MD) trajectories.^5–12^ However, these approaches conflate direct and indirect couplings: two residues may appear linked whenever they are statistically correlated, even if the correlation is spurious (e.g., mediated by a third site).

Bayesian networks (BNs) address this limitation by encoding conditional independence: an edge is retained only when the statistical dependency persists after conditioning on all other variables, yielding a directed acyclic graph (DAG) that encapsulates the non-transitive backbone of the motion.^13–15^ By isolating non-transitive interactions, BN topologies are markedly sparser and more directly interpretable than correlation maps, providing compact, testable hypotheses for downstream mechanistic analysis.

To make this approach broadly accessible, the user-friendly software package, BaNDyT, was developed that enables researchers to construct and analyze Bayesian networks directly from MD trajectory data.^16^ It discretizes per-frame observables, searches-and-scores model space using a minimum uncertainty score to suppress false positives,^13^ and, crucially, treats all frames as exchangeable, producing a frame-independent DAG that captures the ensemble’s overall dependency structure. BaNDyT has been validated on GPCR–G-protein complexes and other systems, demonstrating its ability to recover known allosteric couplings in a broad variety of datasets. This capability forms the foundation for DRUMBEAT’s granular time-resolved analysis.

While BaNDyT generates a robust BN reflecting the entirety of the MD trajectory, it remains agnostic about how residue pair interactions evolve throughout the trajectory. In many systems, timing is critical: for example, WW-domain folding follows parallel pathways distinguished by the order of hairpin pairing;^17^ membrane proteins such as G protein coupled receptors (GPCR) activation proceeds through short-lived intermediate contacts at the ligand pocket and beyond;^18^ and transient salt bridges can gate enzyme catalysis on the microsecond scale.^19^ These observations have motivated dynamic-network representations that attempt to track connectivity over time. Dynamic Network Analysis (DNA) is an example of such a method which provides a time-resolved view of residue interactions by recomputing connectivity patterns throughout a trajectory.^11^ However, approaches such as DNA often suffer from limited interpretability and high noise, as they capture both direct and indirect correlations and can obscure mechanistic insight. There remains a critical need for methods to build on top of BaNDyT and deliver interpretable, temporally resolved maps of cooperative events, enabling clearer mechanistic understanding of biomolecular transitions.

Dynamic Bayesian Networks (DBNs) provide a more rigorous alternative, extending BNs with explicit time-slice nodes and inter-slice edges to model conditional dependencies across frames.^20,21^ While powerful, DBNs suffer from steep practical costs: the state space grows quadratically with the number of variables, parameter learning requires strong priors or very large datasets, and the resulting models are difficult to interpret as causal arrows proliferate both within and between slices. This is exacerbated by the implicit stationarity assumption, which is unrealistic in the protein dynamics context---incorporation of non-stationarity in the DBN framework makes it even more computationally and sample-hungry. Recent applications to protein kinetics therefore impose heavy sparsity constraints or coarse-grain the system into a handful of collective variables, limiting atomistic insight. There remains a need for a method that preserves BN interpretability, scales to thousands of per-frame features, and exposes the temporal evolution of direct dependencies without an exponential increase in parameters - gaps that DRUMBEAT is designed to fill.

DRUMBEAT combines a universal Bayesian network topology with sliding-window rescoring to produce interpretable, temporally resolved maps of cooperative events in molecular dynamics trajectories, enabling the identification of dynamically evolving residue communities and transition events at atomistic resolution.

### Motivation – Dynamically Resolved Network for Temporal Data

Proteins seldom remain locked in one conformation; they shuttle among states, and those transitions often regulate protein function. A static network can capture the dependencies that stabilize a single structure but cannot distinguish those that drive transitions between structures. A frame-independent Bayesian network, such as the one generated by BaNDyT, is excellent at identifying direct couplings over an entire ensemble but is blind to *when* each coupling matters. For folding, ligand binding, or allosteric signaling, the decisive information lies in the brief windows during which new dependencies flare up and old ones are attenuated or disappear. What is needed, therefore, is a model that preserves BaNDyT’s interpretability yet adds an explicit time axis.

DRUMBEAT meets this need via the contiguous rescoring of a universal graph, a single Bayesian network learned once on the concatenated (or stratified-sampled) trajectories (Fig. 1). This fixed topology acts as a common scaffold on which every window of time can be overlayed. Temporal resolution is recovered *post hoc*: a sliding window sweeps through each trajectory, and within each window, we recompute the edge weights, here, via mutual information, without relearning the structure. The two-stage design sidesteps the state-space explosion that plagues Dynamic Bayesian Networks by reassessing direct pairwise dependencies as conditional on the time variable within a specific window. Thus, computational complexity is trivial once the universal graph is recovered, with the procedure easily parallelized. By varying the window size attuned to the protein or protein complex in question, an optimal balance can be achieved between true non-stationarity, resolution, interpretability, and computational efficiency. Table 1 provides a summary of similar methods for direct comparison with DRUMBEAT.

**Figure 1.**
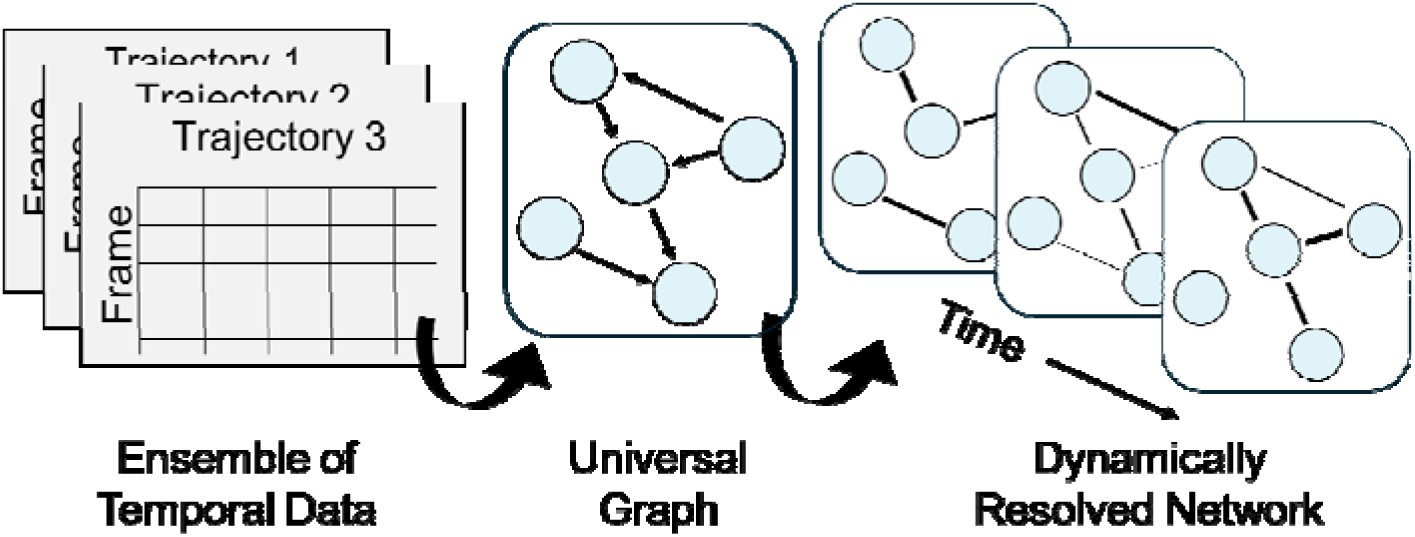
Scheme representing the general dynamic network construction. An ensemble of temporal data is taken as input with the data combined using concatenation or sampling. The resulting dataset is then used to construct a universal Bayesian network graph. Finally, the universal graph topology is fixed, and the edge weights are recomputed over time, providing a dynamic representation of the network for each trajectory.

**Table 1.** A comparative list of data analysis methods that can be applied to Molecular Dynamics data, with BaNDyT and DRUMBEAT included. For each method, a brief summary is provided as well as its possible mechanistic insight.

| Method | Temporally Resolved | Interpretability | Scalability | Brief Summary | Mechanistic Insight Provided |
| --- | --- | --- | --- | --- | --- |
| BaNDyT | No | High | High | Recovers Bayesian networks from entire MD trajectories. | Identifies direct residue dependencies and allosteric couplings in an entire MD trajectory; lacks temporal resolution. |
| DRUMBEAT | Yes | High | High | Dynamically rescores universal Bayesian network graph with sliding-window over MD trajectories. | Identifies direct, time-resolved dependencies and dynamic residue communities; reveals order and timing of transitions |
| Dynamic Network Analysis (DNA) | No | Moderate | High | Recovers dynamic correlation/contact networks. | Captures correlated motions and transient networks but mixes direct/indirect effects; limited mechanistic specificity. |
| Time-lagged independent component analysis (TICA) | No | Moderate | High | Finds slowest collective motions via time-lagged correlations. | Reveals dominant slow modes; highlights collective transitions, but with less direct residue-level mechanistic detail. |
| Dynamic Mode Decomposition (DMD) | Yes | Moderate | Moderate | Decomposes time series into spatial modes and temporal dynamics. | Identifies dominant dynamical modes; mechanistic interpretation can be challenging for complex biomolecular systems. |
| Dynamic Bayesian Network (DBN) | Yes, but only under a stationarity assumption for computationally feasible applications | Low to Moderate | Low | Extends Bayesian networks with explicit time-slice and inter-slice dependencies. | Models conditional independencies across time; interpretability and scalability sharply decrease with system size/complexity. |

Window-specific edge weights carry mechanistic meaning. An edge that is marginal most of the time, but spikes briefly flags a cooperative event; for example, two distant segments forming a contact during transition to a new state. Many graph-level metrics can be mined from these time-varying edges; here we focus on the weighted degree – the sum of edge weights incident on a node in each window. This produces a smooth trace we term a *Time-Resolved Allosteric Community* (TRAC) signal, which pinpoints transition moments with high sensitivity. Other choices (network entropy or betweenness) are also available within DRUMBEAT, and may provide insight on signaling pathways or community reorganization. Together, the universal-graph concept and its dynamic rescoring give us a compact yet powerful lens through which to identify how macromolecules switch, shuffle, and communicate in real time.

The next section details DRUMBEAT application using a representative example of the folding dynamics of the WW domain FiP35. As summarized in the pseudocode in Figure 2, the workflow proceeds in four stages: (i) load an ensemble of trajectories and apply a mutual-information filter to winnow features; (ii) assemble a universal dataset by concatenation or stratified sampling; (iii) learn a universal Bayesian network graph with BaNDyT; and (iv) rescore that fixed topology alongside each trajectory to obtain a time-resolved network representation.

**Figure 2.**
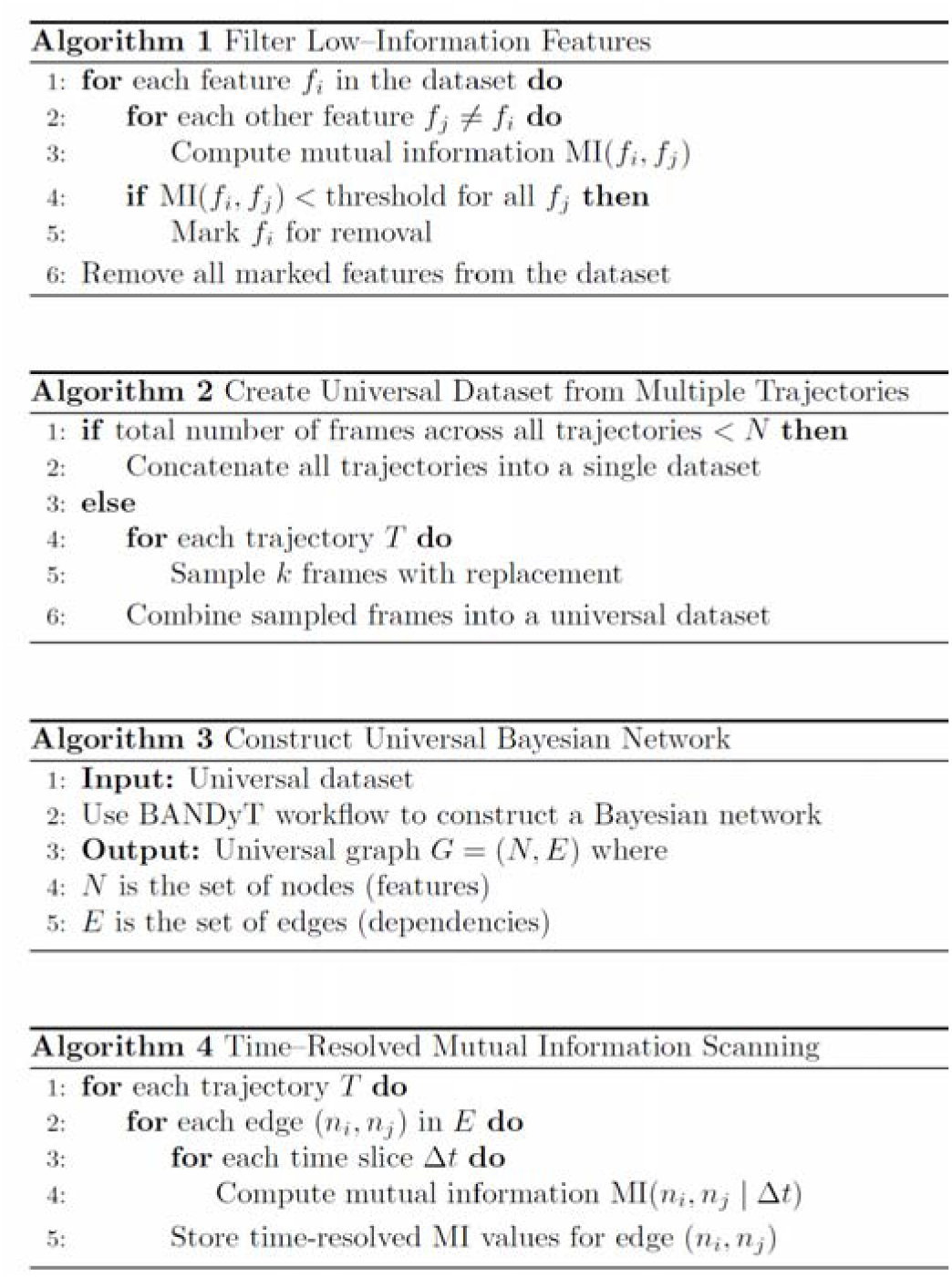
Pseudocode representation for the time-resolved network analysis in DRUMBEAT.

## Methods

### 1. Protein dynamics data

DRUMBEAT is designed for MD trajectories that contain, or may conceal, conformational transitions. In practice, any all-atom or coarse-grained simulation of a protein (or protein complex) can be analyzed: if a large-scale switch is already known, DRUMBEAT will detect it; if not, the algorithm will still expose any state changes that occur along the trajectory.

In the following sections, we describe the usage of DRUMBEAT on FiP35 folding dynamics obtained by D.E. Shaw research group.^17^ The data comprises two 100 μs trajectories with multiple folding events in each trajectory. We aim to utilize DRUMBEAT to (i)confirm the folding transitions and validate the duel-pathway mechanism by which folding is occurring, and (ii) provide new insight into residues involved in folding and granular ordering of events that recapitulate the folding mechanism.

### 2. Preparation of data

Once the raw MD trajectories are available, the first task is to translate each coordinate file into a frame-by-feature matrix that DRUMBEAT can take as input. The user begins by deciding which descriptors best capture the motions of interest. In this work, we focus on binary residue–residue contacts produced by GetContacts^22^(https://www.github.com/getcontacts), but any per-frame quantity (interaction energies, backbone or side-chain dihedrals) can serve as long as it can be organized with frames as rows and features as columns. For contact fingerprints, GetContacts reads the original trajectory files (e.g. xtc or.dcd) and outputs a tab-separated table ready for DRUMBEAT; for other feature types, the user collates the values into a comma-separated sheet (*.csv) of identical dimensionality, one file per trajectory.

Each of the GetContacts files (or user specified.csv files) is loaded directly into DRUMBEAT where the files are converted into Trajectory objects. The resulting trajectory objects will be used downstream for feature filtering, construction of the universal dataset, and ultimately the dynamic network analysis.

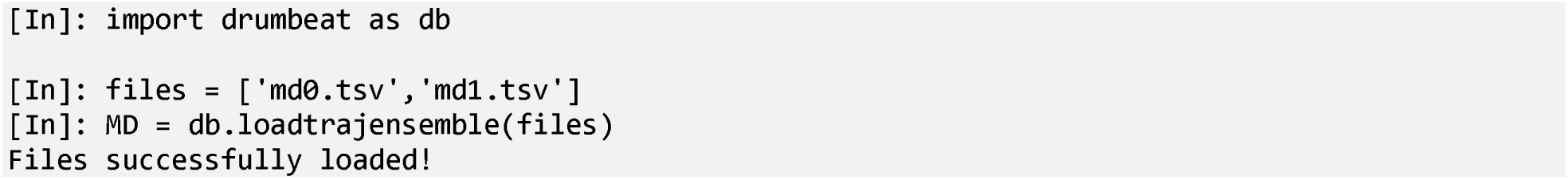

If utilizing.csv files, the user should make sure that the format of the input files is correct. Specifically, DRUMBEAT expects the.csv to have trajectory frames as the rows and features as the columns, with the first row being the feature names.

### 3. Feature selection via Mutual Information

Very large feature sets inflate the computational cost of the universal graph search-and-score, so a light first-pass filter is strongly recommended. DRUMBEAT provides a pairwise mutual information (MI) screen to maximize efficiency given that the computation of the universal graph in the following step scales exponentially. In short, DRUMBEAT creates a pairwise MI matrix for all the features and omits those that fail to have MI with any other contact above a given threshold. We suggest the user proceed in one of two ways. First, an explicit threshold can be used for filtering. The default is 0.005 bits, but the user can use a threshold as high as 0.01 bits, which, in our experience, will still yield the most informative features and not risk losing key information. Second, the user can determine a set number of features they want to work with and utilize the threshold that yields that number of features. In the example case, we utilized a threshold of 0.035 bits, which yielded 198 and 202 contacts for each of the two trajectories, respectively. In our experience, having around 200 features provides a very feasible computation of the universal graph that can be completed on most desktop computers while also capturing the most informative features for the system.

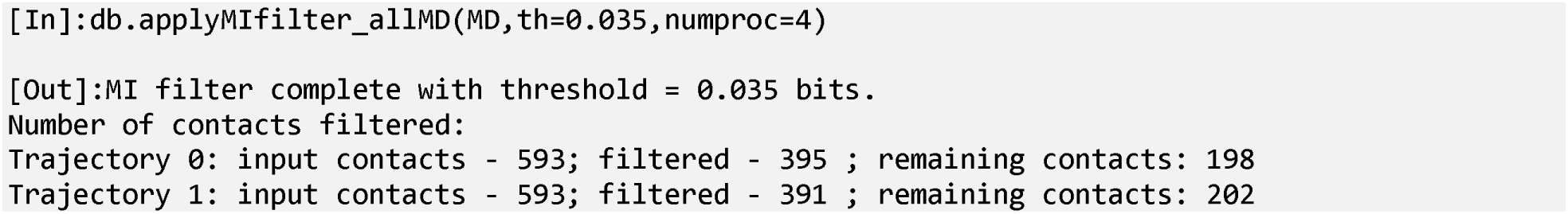

Depending on the size of the trajectory datasets, this computation is moderately time-consuming (in the example above, 15 minutes with 4 CPUs). As shown in the code above, the user is free to utilize more processors to speed up the calculation. Once this step has been completed, DRUMBEAT saves the MI matrix within each object, and the filtering can be done again with a new threshold at a much faster speed.

### 4. Building a universal dataset

With the per-trajectory matrices prepared, the next step is to fuse them into a single universal dataset that captures the entire conformational ensemble. In DRUMBEAT, the default is to first harmonize feature order: each column must refer to the same contact or descriptor in every file, so we start with the intersection of all feature lists and reorder columns identically across trajectories. This incorporates features in 100% of trajectories. Alternatively, if the number of contacts is small, the user may select features that appear in a smaller set of trajectories (i.e., 70%). If the number of features is around 200 (given the step above) and the sum of all trajectory frames is still modest (roughly 10^5^ frames), the user can simply concatenate the matrices end-to-end. This preserves every data point and allows DRUMBEAT to estimate edge strengths with maximum statistical power without additional bookkeeping.

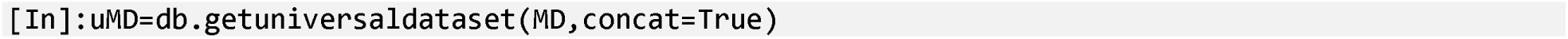

As the ensemble approaches a larger number of features (>200) and longer run times, building the universal graph using a concatenated dataset becomes computationally infeasible. In those cases, we recommend a subsampling scheme where DRUMBEAT uniformly samples, with replacement, frames from each trajectory. This can be achieved by using the same function as above but instead providing the sample size to take from each trajectory as well as a random seed.

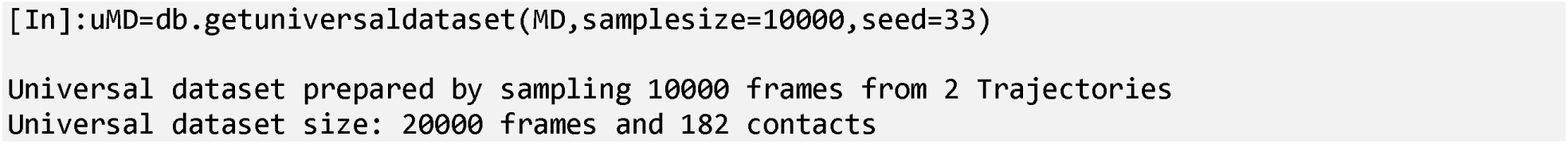

The resulting dataset—20,000 rows by 182 features in this example—strikes a practical balance between statistical power and the runtime of the forthcoming Bayesian network search-and-score.

### 5. Recovering a universal BN graph

With the universal dataset in hand, we next construct a universal BN graph that encodes all direct probabilistic dependencies present across all trajectories and time points. Construction is carried out with BaNDyT, the Bayesian network modeling software previously introduced for frame-independent MD analysis and available open-source at github.com/bandyt-group/bandyt.^14^ Briefly, the universal frame × feature matrix built in step four is passed to BaNDyT, and a Bayesian network is constructed and provided as an output (see Ref. 6 for further detail). We will use the resulting BN primarily as a topology-only object, or DAG, which we hereafter call the universal graph, even though the BN contains edge strength information as well. Figure 3 shows the resulting universal graph for the FiP35 system with the four highest degree nodes highlighted in blue. The four high degree contacts are shown in the Fip35 protein structure in Figure 3B, highlighting their role in both hairpins. Because the graph is learned on pooled snapshots, it is agnostic to when a dependency manifests; temporal resolution is recovered later during DRUMBEAT’s sliding-window rescoring (Step 6).

**Figure 3.**
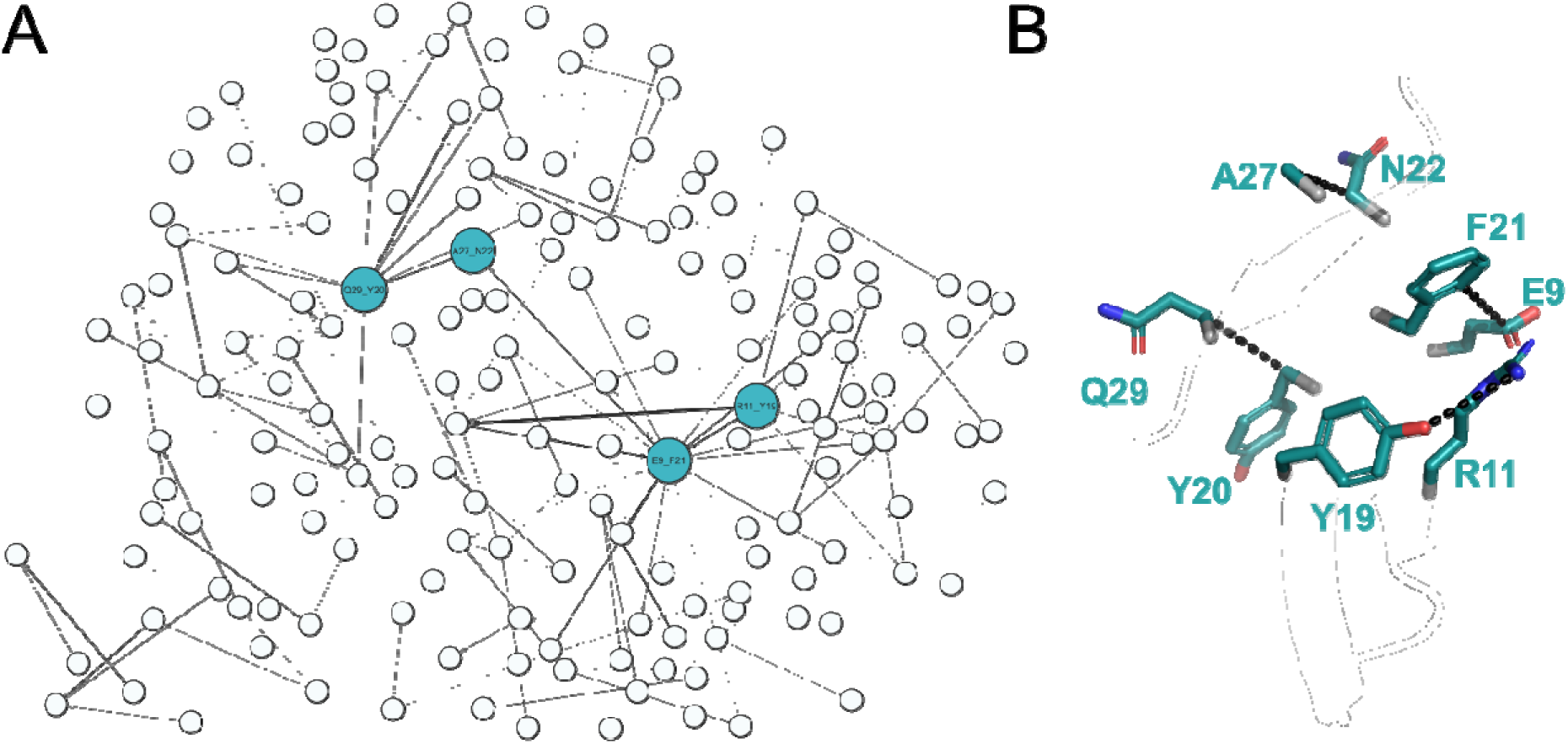
(A) Universal BN Graph built using the BaNDyT software. Four nodes, highlighted in blue, are the highest degree nodes in the universal graph. Edges are shown in varying transparency based on their weight. (B) Fip35 protein with high degree contacts labeled. Contacts span both hairpins, emphasizing the role in the dual pathway mechanism.

The user should be aware that if the number of contacts is large (>200), the expected run times are on the order of hours to days. In the sample case above, we built the universal graph for the universal dataset that has 182 contacts and 20000 frames. The BaNDyT calculation for this dataset took about seven hours on a high-performance computer with a single CPU. For reference, we also built the universal graph for a larger universal dataset (467 contacts and 20000 frames), which took four days on the same hardware. This falls in line with the expectations (BN structure recovery being generally NP-hard) and previous reports that Bayesian network computation time via BaNDyT scales exponentially with the number of features.^14,15,23^

### 6. Rescoring the universal graph alongside the trajectories

At this stage, the universal graph encodes the direct dependencies present across all trajectories and timepoints. The next goal is to determine when, along each trajectory, these dependencies are most salient. DRUMBEAT accomplishes this by scanning each trajectory using the universal graph as a fixed topology, recomputing edge weights within sliding time windows.

The DRUMBEAT rescoring step takes as input the universal graph (generated by BaNDyT in Step 5) and the ensemble of trajectory data.

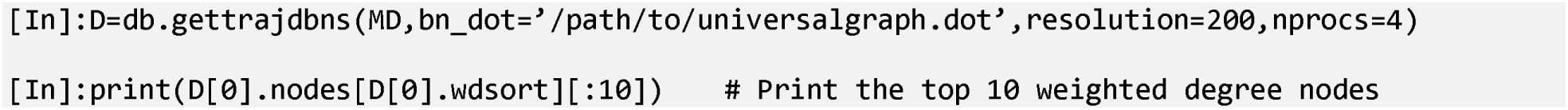

The output is a dynamic network object for each trajectory. For each window along the trajectory, DRUMBEAT recalculates edge weights (using mutual information by default) and derives node-level metrics such as the temporally resolved weighted degree. DRUMBEAT also offers the calculation of other temporally resolved network-level metrics such as entropy. Figure S1 shows the time-resolved entropy for this system. This measure can act as a first pass computation to identify where transitions occur during the trajectory (large changes in entropy correspond to folding transitions; See Figure S1). On the other hand, weighted degree allows the user to identify the specific features involved in those transitions as well as the distinct ordering of events that may enable or facilitate the transition. We discuss this in more detail in the Interpretation section below.

The temporal resolution of DRUMBEAT’s output is determined by the window size, which should be chosen based on the timescale of the transitions of interest and the available computational resources. High temporal resolution (small window size) maximizes detail but can increase noise and computational cost. Conversely, lower resolution (larger window size or down sampling) reduces noise and highlights broader trends, although very large windows may obscure the order of events.

Figure 4 illustrates the effect of window size on the time-resolved weighted degree for the top five nodes. Here we implemented a standard approach for identifying the top five nodes: take the maximum weighted degree value during the trajectory and rank the nodes accordingly (as shown in sample code above). In principle, other criteria can be used, such as focusing on peaks near or before a specific transition. At the highest resolution (Figure 4-top, window size 40 ns; data sampled every 200 ps), the signal is detailed but noisy. Intermediate resolutions, such as 400 ns or 800 ns, provide a balance between signal clarity and temporal detail, allowing transitions and their order to be resolved. At the lowest resolution (e.g., 4 μs, Figure 4-bottom), transitions remain detectable but fine mechanistic details are lost.

**Figure 4.**
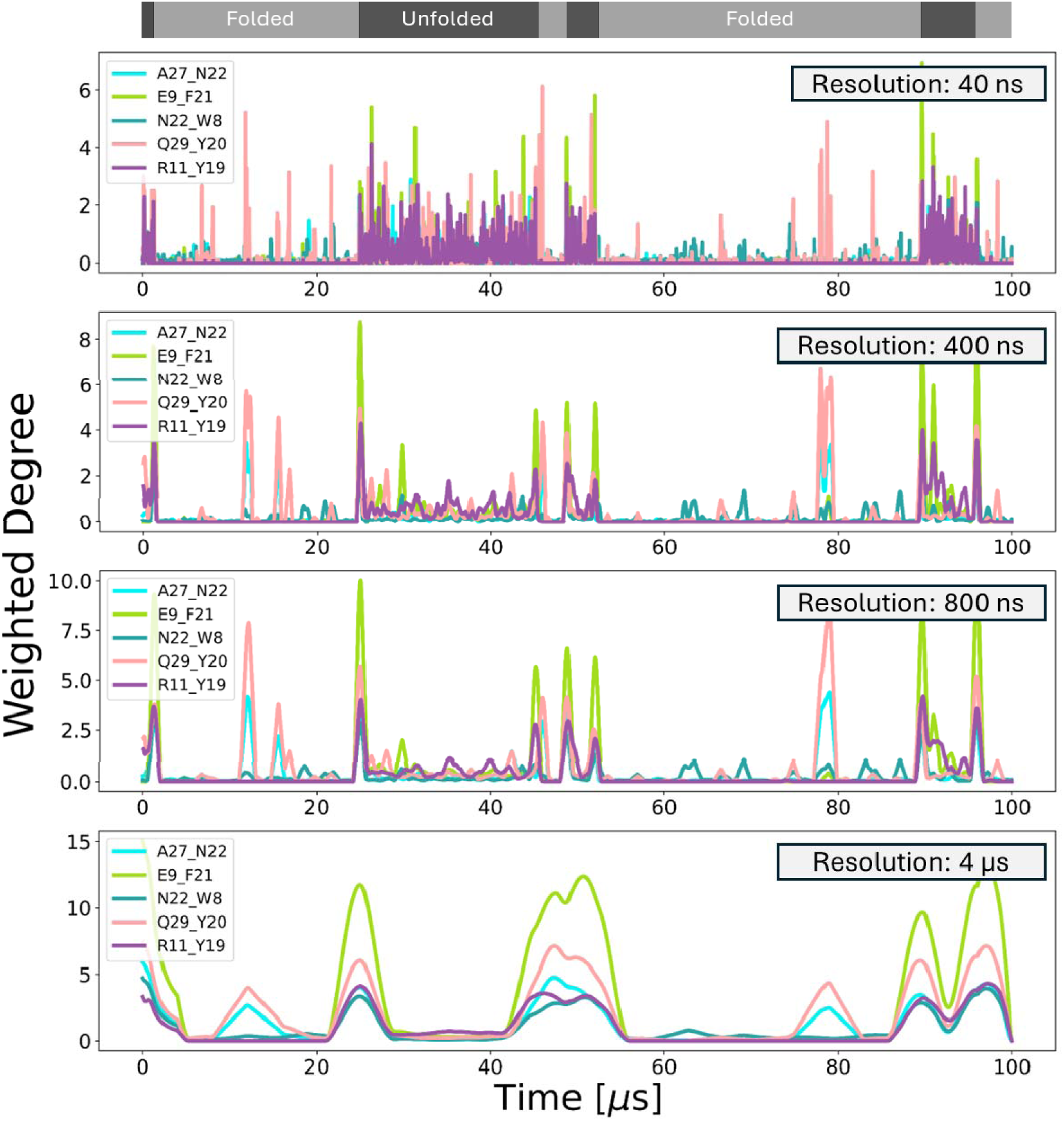
Temporally resolved weighted degree for the top five nodes from DRUMBEAT. The weighted degree is computed by using dynamic network rescoring with MI-based edge weights. Top five nodes are determined by ranking based on max value of temporally resolved weighted degree. At the top are folded and unfolded states as determined by Shaw. Different temporal resolutions and time windows are shown, with the top panel at the highest resolution (40 ns) and the bottom panel at the lowest resolution (4μs). We found that for the FiP35 system, the 400ns resolution showed the best balance between signal and noise.

For the Fip35 system, we found that a window size of 400 ns offers an optimal trade-off, enabling identification of both the timing and sequence of folding events. This approach allows DRUMBEAT to deliver interpretable, granular insights into the dynamics of complex molecular systems.

### 7. Extracting subnetworks for nodes of interest

At this stage, DRUMBEAT provides dynamic network outputs for each trajectory, representing the evolution of network connectivity across time. Users can extract subnetworks and analyze the state space based on nodes of particular interest. The prioritization of nodes for such analyses can proceed in two ways: (1) user-defined nodes of interest, or (2) top nodes identified from DRUMBEAT outputs, such as those highlighted in Step 6. In either scenario, a subnetwork corresponding to the selected nodes can be extracted from the universal graph.

Following the approach outlined above, we concentrated on the top five nodes identified by time-resolved weighted degree. Figure 5A illustrates the subnetwork centered on these five nodes. Specifically, the subnetwork comprises the first-degree probabilistic neighborhood of these nodes, including both the nodes themselves and all their immediate neighbors in the universal graph. The highlighted nodes display a high degree of connectivity, consistent with their importance as determined by weighted degree. Users may further investigate nodes that connect major hubs, or connector nodes, as they may provide further mechanistic insight.

**Figure 5.**
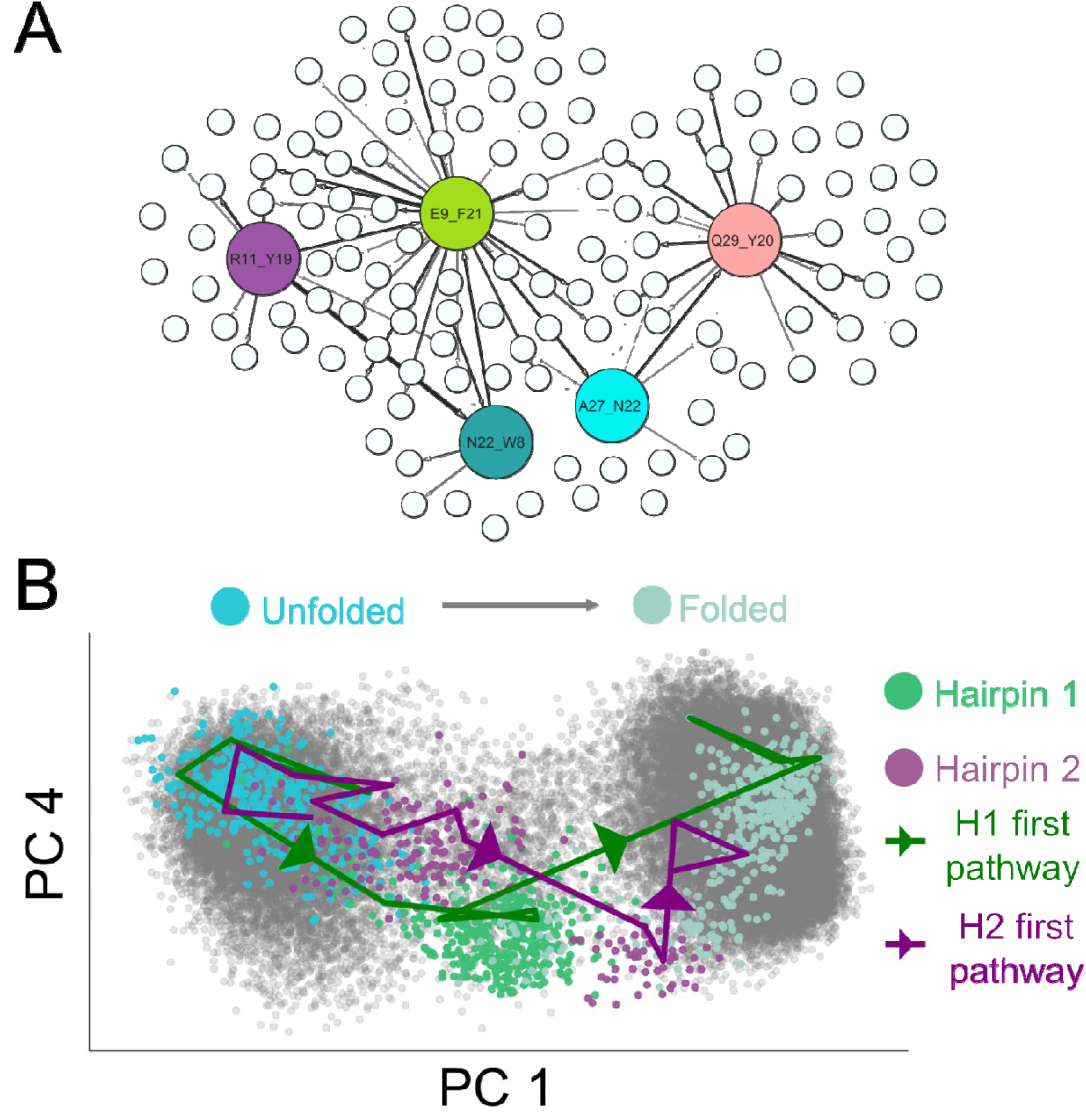
Two examples of analysis output from DRUMBEAT. (A) Subnetwork of key nodes identified by weighted degree analysis. In principle, this can be accomplished for any set of nodes using internal functionalities in DRUMBEAT. (B) Principal component analysis of trajectory data using subset of nodes defined by subnetwork in A. Specific points during folding process are highlighted and showcase two pathways: Hairpin 1 (H1)-first pathway in green and Hairpin 2 (H2)-first pathway in purple.

Although not carried out here, this was demonstrated in a previous report utilizing DRUMBEAT.^24^ Additionally, the entire set of nodes within the subnetwork, including hub nodes and their neighborhoods, can serve as input for further mechanistic modeling and analyses.

As reported by Shaw et al., the FiP35 protein trajectories contain both folded and unfolded states, connected by two principal folding pathways.^17^ DRUMBEAT facilitates further analysis by enabling principal component analysis (PCA) on a subset of nodes, such as those in the subnetwork shown in Figure 5A. Using this subset, DRUMBEAT performs PCA on the entire trajectory (grey points). The resulting projection, shown in Figure 5B, reveals two distinct states in principal component space: unfolded (left) and folded (right).

Further, DRUMBEAT allows for the investigation of the two folding pathways. The first pathway, depicted as a green line, transitions from unfolded to folded by passing through the region defined by the formation of the first hairpin (green points). The second pathway, shown as a purple line, also begins in the unfolded state but traverses the region associated with hairpin 2 formation (purple points). Ultimately, DRUMBEAT’s temporal resolution allows users to identify distinct states, possible intermediates, and pathways connecting these states.

The procedures described in Steps 6 and 7 show that while the core function of DRUMBEAT is to generate a dynamic network from molecular dynamics trajectories, the methodology is designed to support a broad complement of downstream analyses. These final steps represent the secondary, analysis-oriented component of DRUMBEAT: once the dynamic network (the primary output from Step 6) is obtained, users can leverage built-in functions within DRUMBEAT to extract subnetworks, perform dimensionality reduction, or conduct other mechanistic investigations. This flexible analysis framework enables users to tailor their exploration to specific scientific questions, extending the utility of DRUMBEAT well beyond initial network construction.

## Results

### Robustness of the methodology

To assess the robustness of the DRUMBEAT methodology, we evaluated the sensitivity of both the universal graph and subsequent time-resolved results to the sampling procedure used in constructing the universal dataset (see Step 3 above). Recall that, because the size of the trajectory data precludes simple concatenation, we employ a sampling approach to generate the universal dataset. This raises the question of whether the resulting universal graph and downstream analyses are dependent on the particular frames selected during sampling. To address this, we repeated the sampling procedure 100 times, generating 100 independent universal datasets. For each dataset, we constructed a universal graph, which was then used for subsequent time-resolved DRUMBEAT analysis.

First, we assessed the consistency of the universal graphs by comparing the degree of the most highly connected nodes across all replicates. As shown in Figure 6A, the standard deviation in node degree among the top-ranking nodes is minimal, indicating that the most connected, and thus most functionally relevant, regions of the network are highly conserved across different samples. This suggests that the overall topology of the universal graph is robust with respect to the sampling process.

**Figure 6.**
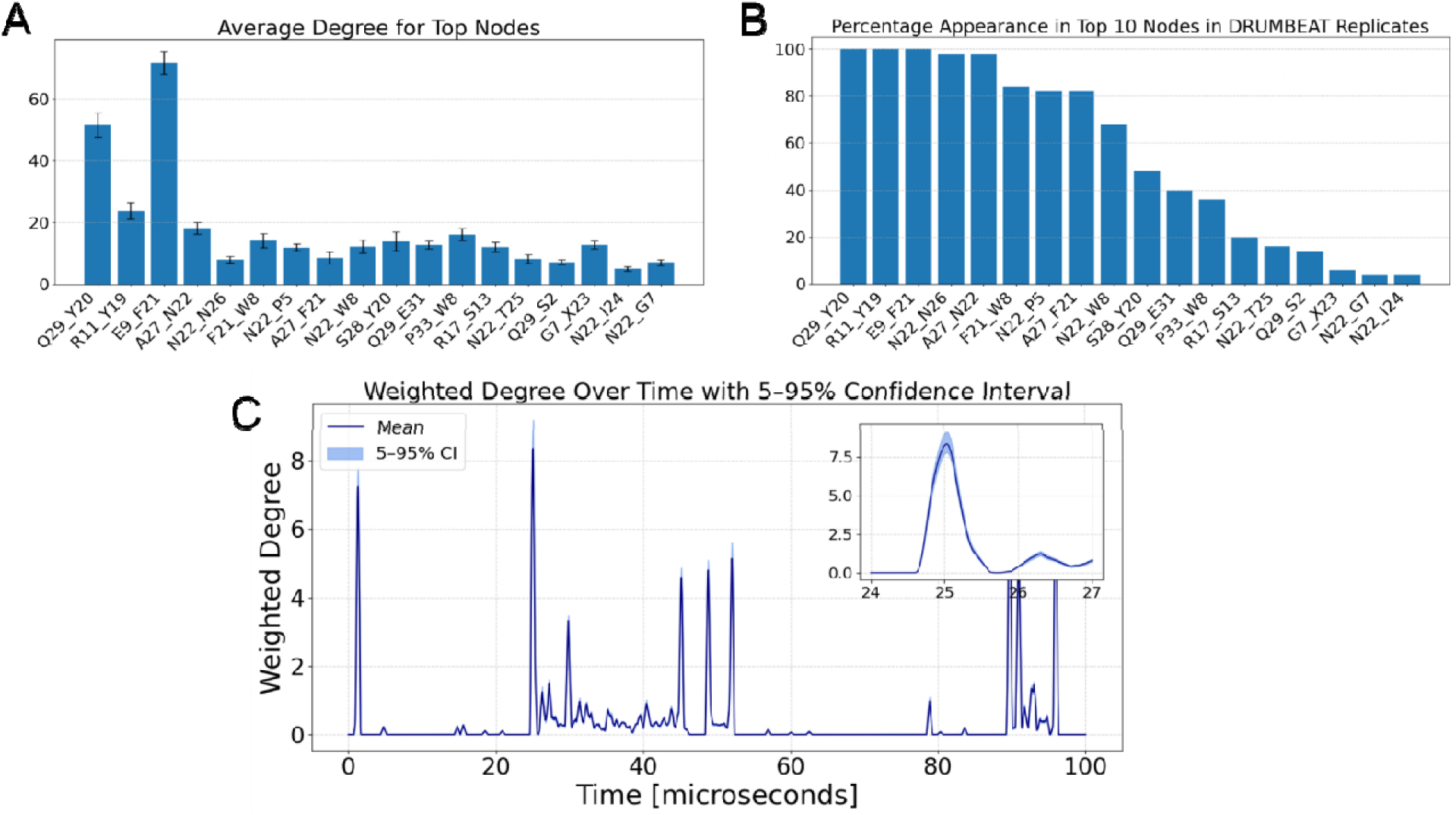
Results from resampling procedure to check for robustness of DRUMBEAT. (A) Average degree for most highly connected nodes in 100 replicas. (B) Appearance of nodes in the top ten weighted degree nodes in 100 DRUMBEAT replica scans. (C) Average temporally resolved weighted degree for E9_F21 node over 100 replicas with 5-95% confidence interval shown with shaded region.

To further test robustness, at the stage of time-resolved analysis, we performed DRUMBEAT scans using each of the 100 universal graphs and extracted the top 10 nodes by weighted degree for each sample. As depicted in Figure 6B, a core set of five nodes appeared in over 95% of the samples, with three of these nodes present in all 100 replicates. This demonstrates that the identification of key nodes by DRUMBEAT is highly consistent and not sensitive to a particular sampling instance.

As an additional check, we examined the time-resolved weighted degree trace for a representative node (E9_F21) across all replicates. The mean and confidence interval for the time-resolved weighted degree are shown in Figure 6C. The variation in this metric is remarkably small, even at peak transition points (see the inset of Figure 6C), indicating that the temporal profiles produced by DRUMBEAT are stable with respect to sampling. These results provide strong evidence that DRUMBEAT’s identification of key nodes and their time-resolved behavior is robust to the specific sampling of trajectory frames.

### Interpretation of DRUMBEAT results

The original Shaw et al. study^17^ provided a landmark dataset for protein folding by performing long-timescale (up to 100 μs) molecular dynamics simulations of the Fip35 WW domain. These simulations captured multiple folding and unfolding events, enabling direct observation of folding mechanisms at atomic resolution. Notably, Shaw and colleagues confirmed that FiP35 folds via two distinct pathways (See Figure 7A): one in which hairpin 1 (formed by the first and second β-strands) closes first, followed by hairpin 2 (second and third β-strands), and another where the order is reversed. Both pathways were observed in their published trajectories. Additionally, Shaw et al. identified specific residues (W8, R11, and F21) as critical for the folding process, based on high ψ-values (ψ measures the resultant change in folding due to mutation)^25^ derived from simulation and validated by experimental data.^17^

**Figure 7.**
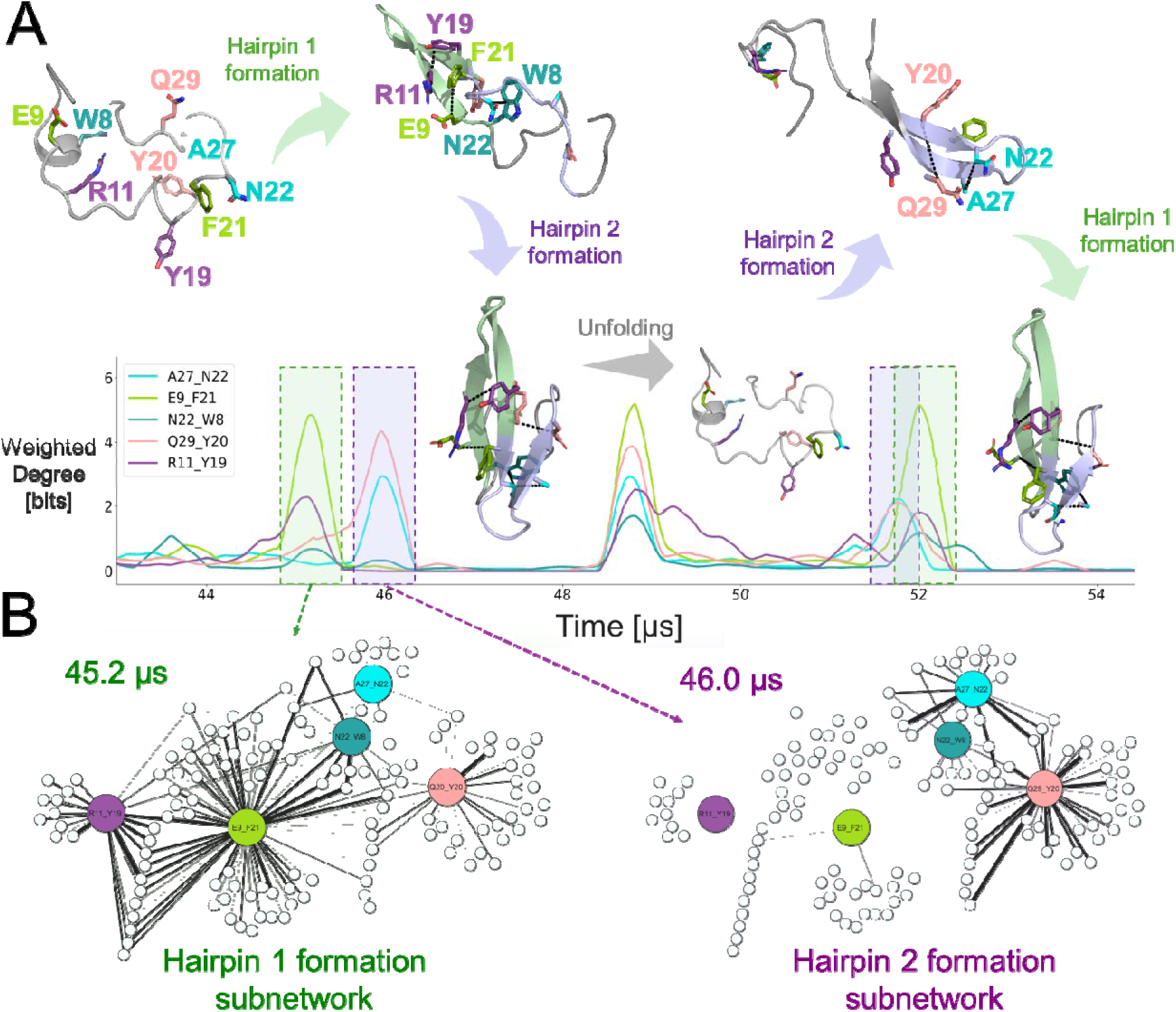
Mechanistic analysis of dual-pathway folding of Fip35 using DRUMBEAT. (A) On the bottom is the plot of the temporally resolved weighted degree for the top five nodes computed for the 100 μs trajectory of FiP35 folding and unfolding. Starting in the unfolded state (top right), the protein initially undergoes the first pathway which has hairpin 1 fold first and the formation of three of five contacts (E9_F21,N22_W8, & R11_Y19), then hairpin 2 fold 1 μs later. Then, the protein unfolds again before refolding via the second pathway. In the second pathway, hairpin 2 forms first resulting in formation of two of five contacts (A27_N22, Q29_Y20), then hairpin 1 forms within 0.5 μs completing the entire folding process. (B) Subnetworks centered around the top five nodes with the temporally resolved edge weights for hairpin 1 & 2 formation at 45.2 μs and 46.0 μs, respectively.

We applied DRUMBEAT to the same Fip35 trajectories to validate these mechanistic findings and to assess whether DRUMBEAT can provide new insights into the folding process. DRUMBEAT identifies key contacts involved in transition events through the detection of temporally resolved allosteric communities (TRACs). As shown in Figure 7, our analysis revealed five principal contacts that are most significant during folding transitions. Of these, three contacts (E9_F21, N22_W8, and R11_Y19) correspond to hairpin 1, while the remaining two (A27_N22 and Q29_Y20) are associated with hairpin 2. Importantly, the residues involved in these contacts include W8, R11, and F21, matching those highlighted by Shaw et al. as critical for folding. This demonstrates that DRUMBEAT, in a fully unsupervised manner, can recover the key residues and contacts previously determined to be underlying protein folding dynamics.

Beyond confirming known results, DRUMBEAT highlights new key residues and provides mechanistic and temporal detail to the analysis. For instance, the contacts E9_F21 and R11_Y19 show how key residues identified by Shaw (F21 & R11) participate in contact formation during folding. Furthermore, the identification of A27_N22 and Q29_Y20 suggests that the closure and stabilization of hairpin 2 play a more prominent, temporally resolved role during certain folding pathways than previously appreciated. Notably, while Shaw et al. (and similar studies analyzing protein contacts) identify which key contacts are involved in the transition, those approaches focus on isolated contacts. In contrast, DRUMBEAT further reveals how these contacts communicate with other regions of the protein allosterically, enabling the analysis of cooperative subnetworks rather than individual interactions. This enhances mechanistic insight into the transition process, uncovering groups of contacts that act together during folding events. By examining the time-resolved weighted degree traces for these contacts (Figure 7A), we can directly observe the sequence of folding events in individual trajectories.

For example, in one trajectory, folding occurs at 45 μs via the first pathway (hairpin 1 closure followed by hairpin 2), as indicated by peaks in the weighted degree of hairpin 1 contacts. In another instance, DRUMBEAT clearly identifies the alternative pathway at 51 μs, where hairpin 2 forms first, followed by hairpin 1. Thus, DRUMBEAT not only pinpoints the critical contacts and residues involved in folding but also provides precise timing and mechanistic context for each folding event.

A key feature of DRUMBEAT is the ability to extract the subnetworks associated with specific transitions. In Figure 7B, we show the subnetworks centered on the top five nodes identified in the DRUMBEAT analysis. The corresponding time-resolved edge weights are shown at the precise moments of hairpin formation (hairpin 1 – 45.2 μs; hairpin 2 – 46.0 μs). During hairpin 1 formation (left panel), edges surrounding the central hairpin 1 nodes and their probabilistic neighbors become prominent, highlighting the coordinated motion underlying the transition. Similarly, during hairpin 2 formation (right panel), edges around the central hairpin 2 nodes are activated, reflecting their involvement in the structural change.

DRUMBEAT offers two major advantages: (1) it identifies key residues and contacts involved in conformational transitions and (2) it provides temporal resolution, revealing the exact timing and sequence of events during transitions. This level of insight is invaluable for understanding the mechanisms of protein folding and could be extended to study other dynamic processes such as allosteric signaling or ligand binding. Moreover, DRUMBEAT’s framework for temporally resolved network analysis could be integrated with other modeling approaches to further dissect the kinetics and pathways of biomolecular transitions.

## Discussion

DRUMBEAT advances the analysis of molecular dynamics (MD) trajectories by enabling unsupervised, time-resolved identification of residue interactions during protein conformational transitions. Applied to the Fip35 WW domain folding dataset from Shaw et al., DRUMBEAT not only recapitulates established folding pathways and critical residues but also uncovers new mechanistic details.

Shaw et al. identified folding pathways and key residues using targeted, prior knowledge-driven analysis and manual curation and inspection. In contrast, DRUMBEAT extracts these features directly from raw trajectory data in an autonomous, fully data-driven manner, without prior knowledge or manual intervention. Furthermore, it enables detailed temporal resolution, revealing the precise order and timing of contact formation events, and identifies additional contacts and residues not previously reported.

Key Features of DRUMBEAT

- Temporally resolved dynamic network modeling
- Straightforward, seamless application to MD trajectories
- Automated detection and prioritization of key residues and contacts
- Automated identification of cooperative residue communities
- Discovery of novel key contacts, accompanied by granular mechanistic interpretation

DRUMBEAT bridges the gap between interpretability and dynamic resolution in MD analysis, providing both confirmation of known mechanisms and discovery of new features in an efficient, fully automated manner.

## Data Availability

The Fip35 simulations used for this work was acquired with permission from DE Shaw Research. Detailed information for the simulations can be found in the published work by Shaw et al.

## Code Availability

DRUMBEAT source code is available on GitHub at (https://github.com/bandyt-group/drumbeat).

## Acknowledgements

This work was funded by grants from the National Institutes of Health R01-GM117923, R35-GM156498 to NV, R01-LM013876 to NV, AR, SBh, and SBr, and R01-LM013138 to ASR. The content is solely the responsibility of the authors and does not necessarily represent the official views of the National Institutes of Health. Additional support is acknowledged by Dr. Susumu Ohno Endowed Chair in Theoretical Biology (held by ASR), and Susumu Ohno Distinguished Investigator Fellowship (to GG). This work was supported in part through the City of Hope’s High Performance Computing resources, services, and staff expertise provided under the ETG. Additionally, we thank DE Shaw Research for providing Fip35 folding trajectories.

## Author contributions

BM developed the methodology, acquired and curated data, implemented and applied the DRUMBEAT software, interpreted the results of the analysis, and wrote the first draft of the manuscript. EM interpreted DRUMBEAT analysis results and developed BaNDyT. GG developed BNOmics generalist BN modeling software utilized by BaNDyT and DRUMBEAT. BM, EM, SBh, NV, ASR, SBr conceptualized the study, and revised and edited the final version of the manuscript. SBh, NV, ASR, SBr acquired funding and supervised the study.

## Competing interests

The authors declare no competing interests.

## Materials & Correspondence

Correspondence and materials requests should be addressed to BM at, SBh at, NV at, ASR at, or SBr at.

## Notes

https://github.com/bandyt-group/drumbeat

### Competing Interest Statement

The authors have declared no competing interest.

