## Supplementary Materials for "DRUMBEAT: Temporally resolved interpretable machine learning model for characterizing state transitions in protein dynamics"

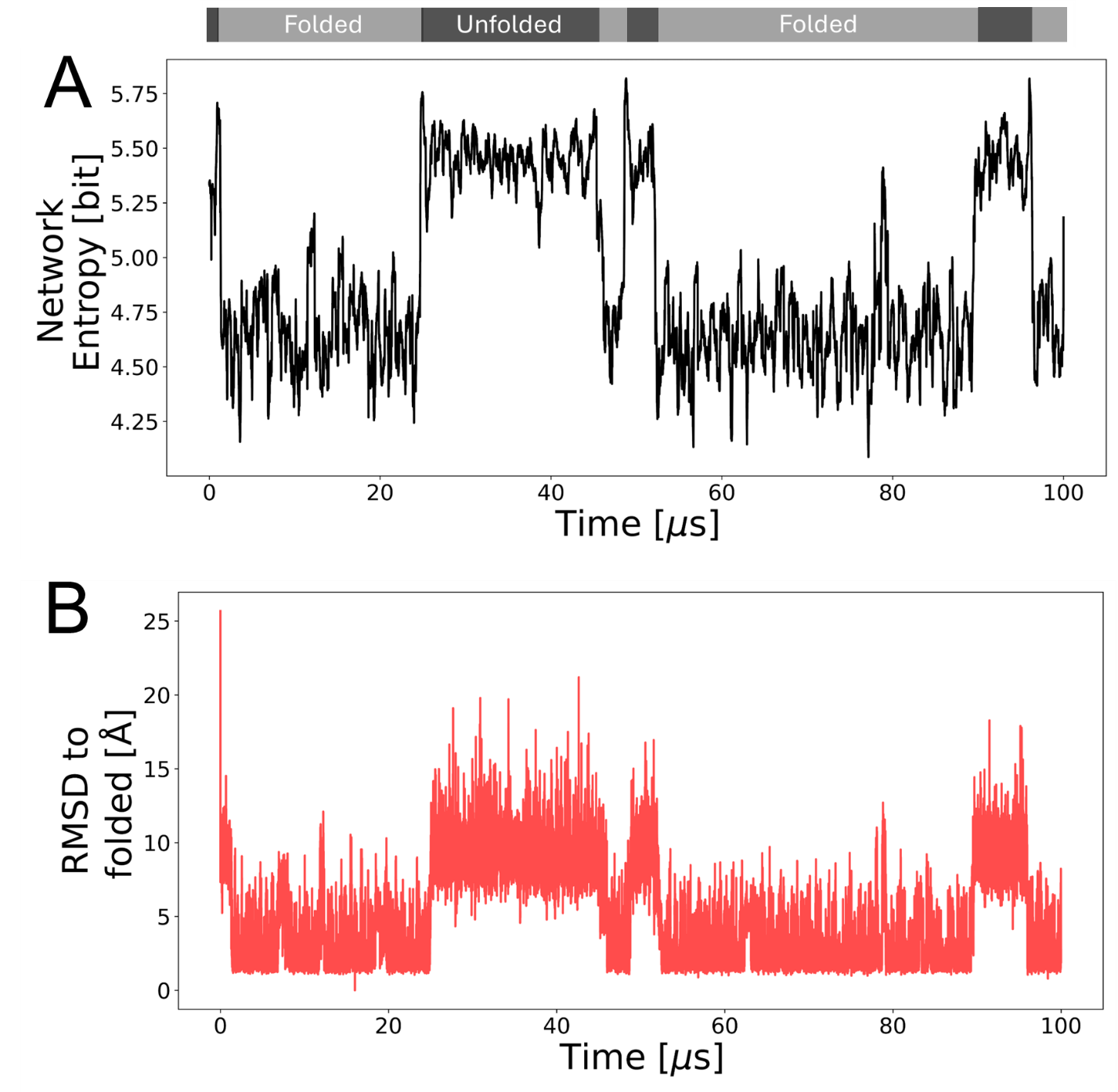

**Figure S1** (A) Network entropy computed over time using sliding window method from DRUMBEAT. (B) RMSD to folded protein state over trajectory. The measurement in B was used by Shaw to report changes between folded and unfolded states. In A we compute the network entropy and show significant changes corresponding to state changes. At the top folded and unfolded states are shown using black (unfolded) and grey (folded) squares as given by Shaw et al.
